# Habitat complexity in organic olive orchards modulates the abundance of natural enemies but not the attraction to plant species

**DOI:** 10.1101/2021.02.04.429588

**Authors:** Hugo Alejandro Álvarez, Marina Morente, Francisca Ruano

**Author notes:** Corresponding author at: Department of Zoology, Faculty of Sciences, University of Granada, Av. Fuente Nueva s/n 18071. Granada, Spain. *E-mail addresses*, (H.A. Álvarez).

## Abstract

Semi-natural habitat complexity and organic management could affect the abundance and diversity of natural enemies and pollinators in olive orchards. Nonetheless, in such agroecosystems the effect of plant structure, plant richness, and plant attraction on the arthropod fauna has been poorly documented. Here we evaluate the influence of those effects jointly as an expression of arthropod abundance and richness in olive trees, ground cover, and adjacent vegetation within organic olive orchards. For this, we used generalized linear models and non-metric multidimensional scaling (NMDS) integrating generalized additive models. Our results suggest that natural enemies and pollinators are mainly attracted to *A. radiatus*, *D. catholica*, and *L. longirrostris* within ground cover and *G. cinerea speciosa*, *Q. rotundifolia*, *R. officinalis*, *T. zygis gracilis*, and *U. parviflorus* within adjacent vegetation. Accordingly, habitat complexity showed a positive relationship with the abundance of key families of natural enemies and pollinators but not with the number of taxa. NMDS showed that plant richness and plant arrangement and scattering affected the key families differently, suggesting that each key family responds to their individual needs for plant resources but forming groups modulated by complexity. This pattern was especially seeing in predators and omnivores. Our findings support that the higher the plant richness and structure of a semi natural-habitat within an olive orchard, the higher the abundance and richness of a given arthropod community (a pattern found in natural ecosystems). The information presented here can be used by producers and technicians to increase the presence and abundance of natural enemies and pollinators within organic olive orchards, and thus improve the ecosystem services provided by semi-natural habitats.

**Graphical abstract:** 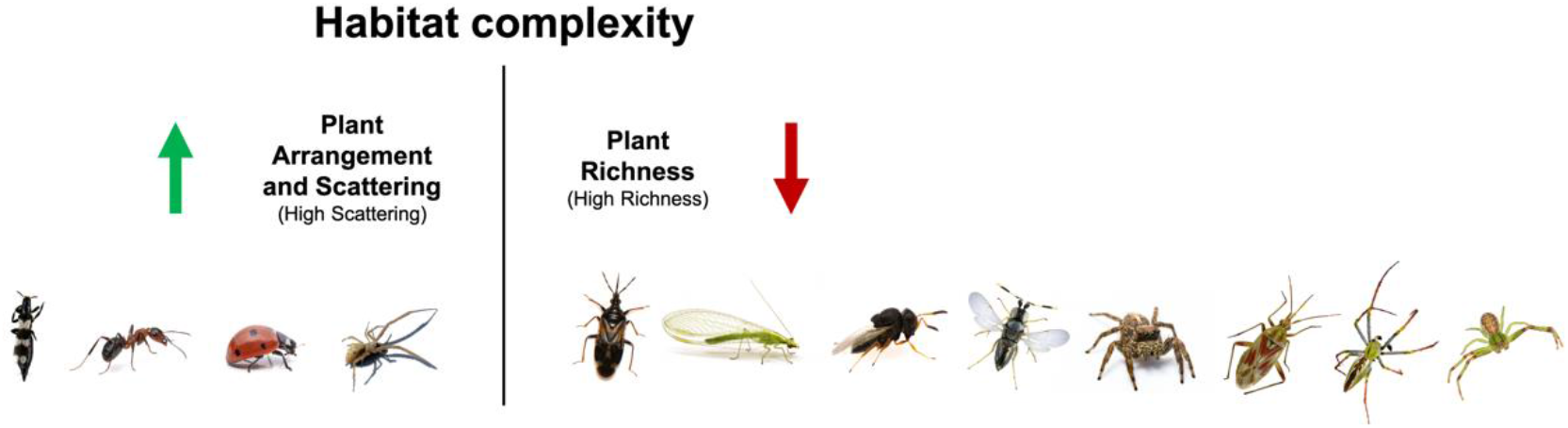

## 1. Introduction

Recent studies have suggested an improvement in the abundance of natural enemies and pollinators due to the presence of semi-natural habitats in agroecosystems (Clemente-Orta and Álvarez, 2019; Karp et al., 2018; Tscharntke et al., 2016). In the European Union policies are currently being implemented with the aim of restoring semi-natural habitats, such as ground cover and adjacent vegetation within vineyards, citrus, almond, and olive orchards (Malavolta and Perdikis, 2018). Especially, there is a growing interest in suitable plant species for ecological restoration and ecosystem services, for example, to prevent soil erosion, maintain soil fertility and control insect pests (biological control) (Oldfield, 2019; Pedrini et al., 2019).

Semi-natural habitat complexity and the management of the ground cover positively affect abundance and variability of natural enemies and pollinators in olive orchards (Álvarez et al., 2019a; 2019b; 2021; Gkisakis et al., 2016; Karamaouna et al., 2019; Villa et al., 2016a). However, a positive or negative response shown by an organism to a nearby habitat could be driven by the structure and composition of such a habitat (Álvarez et al., 2016; 2017; 2021; Balmford et al., 2012; Clemente-Orta et al., 2020;

Laurance, 2007; López-Barrera et al., 2007; Wan et al., 2018). Indeed, the composition and dominance of plant species are key factors in natural habitats that drive the richness and abundance of insects, i.e., the higher the plant species richness in a habitat, the higher the richness and abundance of a given insect community. Interestingly, this pattern is especially reflected on predator insects (Haddad et al., 2001; Knops et al., 1999).

It is known that some plant species can attract more natural enemies and pollinators than others, which is due to the form of functional traits of flowers or prey presence, amongst other factors (Hatt et al., 2017; Nave et al., 2016; Van Rijn and Wackers, 2016). In olive orchards some plant species within ground cover and adjacent vegetation have shown positive effects on predators, omnivores (Torres, 2006), and pollinators (Karamaouna et al., 2019). For example, in a previous study Álvarez et al. (2019a) suggested that arthropod abundance is affected by the type of vegetation, i.e., most plant species within ground cover and adjacent vegetation had a higher abundance of natural enemies than the olive trees, although each trophic guild of natural enemies (e.g., omnivores, parasitoids, and predators) had a specific relationship with a type of vegetation. They showed that four herbaceous species within ground cover attract more predators than others, and six shrubby species within adjacent vegetation attract more omnivores than others. Nevertheless, there is no characterization of the arthropod fauna that is attracted to such plants and no quantification of the effects of that plants on arthropod abundance.

Despite the efforts of different authors to assess the effects of ground cover and adjacent vegetation on natural enemies, pollinators, and olive pests, to the best of our knowledge there is no study that has focused on (1) the attraction of the (whole) arthropod fauna to key plant species within ground cover and adjacent vegetation and (2) the effects of habitat complexity on arthropod attraction. This is of great importance because identifying habitat features and plant species that could attract more key (beneficial) arthropods will be paramount to improve the organic management of olive orchards by means of conserving and planning ground cover and adjacent vegetation, given the fact that, for example, not all natural enemies in a semi-natural habitat are able to produce an effective biological control of pests (Karp et al., 2018; Rusch et al., 2010). Based on the above, the aim of this study was to assess the potential effects of plant species within the ground cover and adjacent vegetation, and the influence of habitat complexity, to attract natural enemies of olive pests and pollinators within organic olive orchards.

## 2. Material and methods

### 2.1. Study area

The study was conducted in organic olive orchards (186.45 ha), in the localities of Píñar (37°24′N, 3°29′W) and Deifontes (37°19′N, 3°34′W), in the province of Granada (southern Spain). The orchards maintained a ground cover for at least three consecutive years. *Bacillus thuringiensis* was used as a preventive pest control for the carpophagous generation of *Prays oleae* (larvae) in July in the orchards of Píñar (for detailed information on climatic conditions and sample areas see Álvarez et al., 2019a). Five different areas with patches and/or hedgerows of adjacent vegetation were sampled within the olive orchards: Deifontes (DEI), Píñar 1 (PI-1), Píñar 2 (PI-2), Píñar 3 (PI-3) and Píñar 5 (PI-5). Based on the soil uses for Andalusia obtained from the information system of occupation of the Spanish soil database at 1:10 000 (SIOSE, www.siose.com) in ArcGis software, and following the definition of classes in the technical guide of the Andalusian soil vegetation cover and uses map (Junta de Andalucía, 2007), we found three soil uses in the sampled areas: (1) woody-sparse scrub: sparse oak (WSS); woody-dense scrub: sparse oak (WDS); and sparse scrub with grass and rocks (SSGR). Accordingly, the structure of the adjacent vegetation was different for each area, i.e., DEI had a surface of WSS and SSGR, PI-1 had a surface of WSS, and PI-2 had a surface of WDS, in contrast PI-3 and PI-5 had one hedgerow (formed primarily by oak trees) and the difference between PI-3 and PI-5 is that the hedgerow in PI-5 is entirely linear (Table 1).

**Table 1.** Plant species richness, arrangement, and scattering in the five study areas of organic olive orchards: Deifontes (DEI), Piñar 1 (PI-1), Piñar 2 (PI-2), Piñar 3 (PI-3), and Piñar 5 (PI-5). Plant species in adjacent vegetation: *C. albidus* (Ca), *G. cinerea speciosa* (Gcs), *P. dulcis* (Pd), *Q. rotundifolia* (Qr), *R. sphaerocarpa* (Rs), *R. officinalis* (Ro), *T. mastichina* (Tm), *T. gracilis* (Tzg), *U. parviflorus* (Up). Plant species in ground cover: *A. radiatus* (Ar), *C. melitenses* (Cm), *D. catholica* (Dc), *E. cicutarium* (Ec), *L. longirrostris* (Ll), *S. vulgaris* (Sv), and the miscellaneous plants (mi).

| Area | Adjacent vegetation |  | Ground cover | Level of richness | Level of arrangement and scattering | Complexity |
| --- | --- | --- | --- | --- | --- | --- |
|  | Richness | Arrangement | Richness |  |  |  |
| DEI | Qr - Rs - Ro - Tzg - Up - Ca - Pd - Tm | Surface: sparse. All plants gathered into clusters | Ll - Dc - Cm - Ec - Sv - mi | 3 | 3 | ++++++ |
| PI-1 | Qr - Rs - Ro - Tzg - Up | Surface: sparse. All plants gathered into clusters | Ll - Dc - Ar - mi | 2 | 3 | ++++ |
| PI-2 | Qr - Rs - Ro - Tzg - Up - Gcs | Surface: dense. All scattered | Ll - Dc - mi | 2 | 4 | ++++++ |
| PI-3 | Qr - Ro | Edge | Ll - mi | 1 | 2 | +++ |
| PI-5 | Qr - Ro | Edge (lineal) | Ll - mi | 1 | 1 | ++ |

### 2.2. Sampling design and habitat classification

We focussed our efforts on collecting samples of arthropods (1) in the most abundant and recognizable (blossom) plant species within adjacent vegetation and ground cover, and (2) in the canopy of the olive trees. Nine plant species were abundant enough in the adjacent vegetation for sampling, i.e., two species of trees: *Prunus dulcis* (Mill.) D.A. Webb, and *Quercus rotundifolia* Lam., and seven species of bushes: *Cistus albidus* L., *Genista cinerea speciosa* Rivar Mart. & al., *Retama sphaerocarpa* (L.) Boiss., *Rosmarinus officinalis* L., *Thymus mastichina* L., *Thymus zygis gracilis* (Boiss.) R. Morales and *Ulex parviflorus* Pourr. Six species of herbaceous plants were abundant enough in the blossom period in the ground cover: *Anacyclus radiatus* Loisel, *Centaurea melitenses* L., *Diplotaxis catholica* (L.) DC., *Erodium cicutarium* (L.) L’Hér, *Leontodon longirrostris* (Finch & P.D. Sell) Talavera, and *Senecio vulgaris* L. In addition, we collected samples in sections of the ground cover located under the canopy of olive trees, i.e., evergreen plants (due to the drip irrigation of the olive trees and their shade) which did not present blooming flowers but formed a distinctive stratum, and thus it was considered as another plant-category named “miscellaneous”. Overall, 17 different plants were measured.

We used as an experimental unit (sample) a suction plot that was a 30 s-suction in a 50 × 50 cm surface of foliage. We used a modified vacuum device CDC Backpack Aspirator G855 (John W. Hock Company, Gainsville, FL, USA) to collect the arthropods. This method allows us to standardize sampling amongst different types of plants (i.e., herbaceous, shrubs, and trees). Samples in this study were collected, weather permitting (once a month) from May to July 2015, which are the months of highest arthropod abundance in olive orchards (Ruano et al., 2004; Santos et al., 2007). We collected 20 randomly distributed samples per plant species in each sample area, depending on plant species availability (see Table 1), and 40 randomly distributed samples in the olive trees. The samples were stored individually and maintained at −20°C until the specimens were classified. The arthropods were identified to family level, unless otherwise specified and classified by trophic guilds, i.e., natural enemies: omnivores, parasitoids, and predators; pollinators and specialist olive pests. The families that were identified as neither natural enemies nor pests were gathered together in a group named neutral arthropods (Wan et al., 2014). Guild classification was based on literature data (see Supporting information in Álvarez et al., 2019a). We pooled together raw sample data by plant species and month.

On the other hand, we followed Alvarez et al. (2019a) to establish a gradient of habitat complexity, however we did not quantify it directly but rather its components, i.e., plant species composition (e.g. plant richness) and habitat structure (e.g. plant arrangement and scattering) for each study area. Table 1 summarizes the different features of each study area and shows the resulting amount of habitat complexity and the level of its components. The level of richness follows the number of plant species in each study area, and then areas were numbered from most to least. The level of arrangement and scattering is based on the information of the structure of the adjacent vegetation given by the SIOSE database (see above). The variables used were (1) surface or hedgerow, where the former is more important, (2) type of soil use: dense or sparse (and their features), where a dense vegetation with woods is more important, and (3) plants scattered across the area or gathered into one or several clusters, where the former is more important. Then the areas were numbered from most to least. We gave the same weight to plant richness and plant arrangement and scattering to express the amount of habitat complexity. Finally, the study areas can be arranged from most to least complex as: PI-2 and DEI (equal), PI-1, PI-3, and PI-5 (Table 1).

### 2.3. Data analysis

Contrary to Álvarez et al. (2019), we analysed the effects of plant-species attraction and habitat complexity on each family of beneficial arthropods. Firstly, to see plant-species effects on arthropod abundance we analysed family abundance per guild (i.e., omnivores, parasitoids, predators, and pollinators) per plant species, for which several generalised linear models (GLMs) were constructed using “quasi-likelihood” with Poisson-like assumptions (quasi-Poisson) tendency (for justification on this approach see Ver Hoef and Boveng, 2007). We fitted models including abundance as the dependent variable and family (per guild) as the independent variable. In this study we considered month samples as independent. If there were significant differences amongst families in a given plant species, the plant species and the most abundant families in that plant were recorded, reported, and expressed in a graphical representation. Nonetheless, the presence and abundance of the total arthropod community by family in plants was still recorded and reported (Appendix).

Secondly, to see habitat effects on arthropod abundance we used the data of the resulting arthropod families that showed the best effects (hereafter called key families), for which four GLMs were constructed using “quasi-likelihood” with Poisson-like assumptions (quasi-Poisson) tendency. Two GLMs were fitted including (1) total abundance of key families and (2) the number of key families as the dependent variables and study area as the independent variable. The next two GLMs were fitted including (1) total abundance of key families and (2) the number of key families as the dependent variables and the level of richness plus the level of arrangement and scattering and their interaction as independent variables. There was no interaction between the two levels, then it was not integrated in further analyses.

Finally, to assess how plant richness and plant arrangement and scattering affect arthropod abundance, the composition of key families and the plants per area were subjected to a non-metric multidimensional scaling (NMDS). Based on the NMDS, smooth surfaces were generated with the data of abundance for each key family. Smooth surfaces result from fitting thin plate splines in two dimensions using generalized additive models. The function selects the degree of smoothing by generalized cross-validation and interpolates the fitted values on the NMDS plot represented by lines ranking in a gradient (Oksanen et al., 2018). Key families were grouped based on the topology of the smooth surfaces in NMDS plots because its form is driven by the maximum abundance showed by plants and study areas. Then, key families were ordered following their relationship with study areas, and therefore, with the level of richness and the level of arrangement and scattering.

All analyses were computed in the R software v.3.6.2 (R Developmental Core Team, 2019). The “vegan” package (Oksanen et al., 2018) was used to compute NMDS and smooth surfaces.

## 3. Results

A total of 9,279 individuals were collected. The arthropods were comprised in 12 orders: Araneae, Blattodea, Coleoptera, Diptera, Dermaptera, Hemiptera, Hymenoptera, Lepidoptera, Mantodea, Neuroptera, Phasmida, Raphidioptera, and Thysanoptera. Overall, 106 families were identified, 51 families were identified as natural enemies and grouped in three trophic guilds: 2 omnivores, 17 parasitoids, and 32 predators. Two families were identified as specialist pests of olive orchards and 1 family as a pollinator. The rest of the families were grouped as neutral arthropods. The Appendix summarizes information of each arthropod family regarding their relative abundance, trophic guild, and records in plants and months. In addition, the plants that showed more abundance and the type of vegetation in which each family was present is detailed.

### 3.1. Difference in arthropod abundance amongst plants

#### 3.1.1. Predators

We found that the abundance amongst families of predators significantly increased in olive trees (*F*_30, 62_ = 6.212, *p* = 0.001), the miscellaneous plants (*F*_30, 62_ = 4.659, *p* = 0.001), and 7 of the 15 plant species sampled, and thus, within the adjacent vegetation abundance was different in *G. cinerea speciosa* (*F*_30, 62_ = 6.537, *p* = 0.001), *Q. rotundifolia* (*F*_30, 62_ = 7.951, *p* = 0.001), *R. officinalis* (*F*_30, 62_ = 7.742, *p* = 0.001), *T. zygis gracilis* (*F*_30, 62_ = 6.786, *p* = 0.001), and *U. parviflorus* (*F*_30, 62_ = 2.956, *p* = 0.001), but within ground cover family abundance was different in *A. radiatus* (*F*_30, 62_ = 4.703, *p* = 0.001) and *D. catholica* (*F*_30, 62_ = 4.804, *p* = 0.001).

Figure 1 graphically summarizes the results of the later analyses and shows the proportion of the families of predators that were significantly more abundant in each plant species. The families of predators that showed best effects on these plants were: Anthocoridae, Miridae (Hemiptera); Oxiopidae, Salticidae, Thomisidae, Uloboridae (Araneae); Aeolothripidae (Thysanoptera); Coccinelidae (Coleoptera); and Chrysopidae (Neuroptera). However, the most representative families of predators were (from most to least abundant): Miridae, Aeolothripidae, and Thomisidae.

**Fig. 1.**
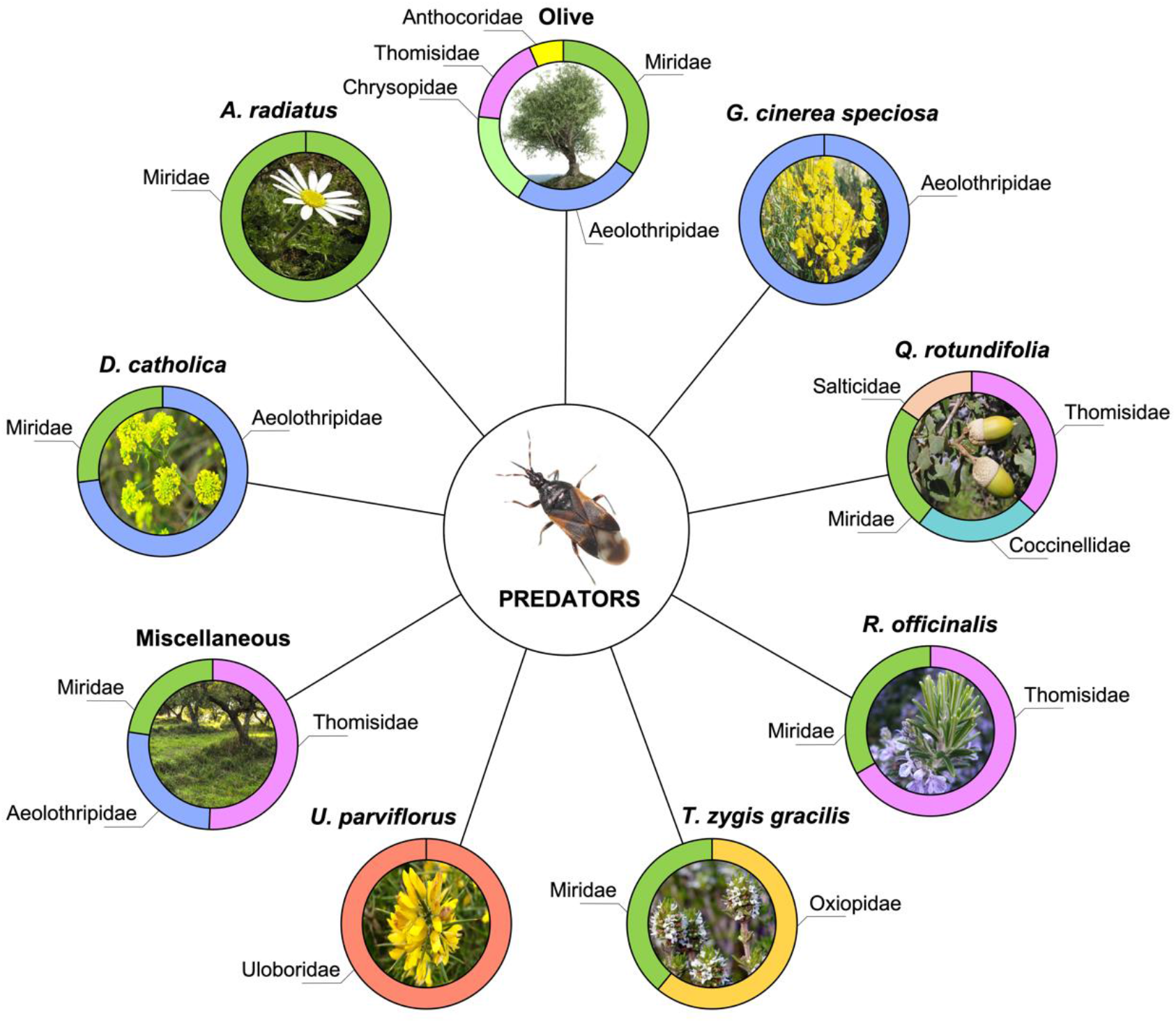
Families of predators that had the highest effects per plant species within organic olive orchards. The circles show the proportion of the abundance amongst such families. Only the plants that showed significant differences on the abundance of predators are shown.

#### 3.1.2. Parasitoids, omnivores, and pollinators

The families of parasitoids were significantly more abundant in olive trees (*F*_13, 28_ = 4.505, *p* = 0.001), with Encyrtidae and Scelionidae being the families that showed best effects (Fig. 2).

**Fig. 2.**
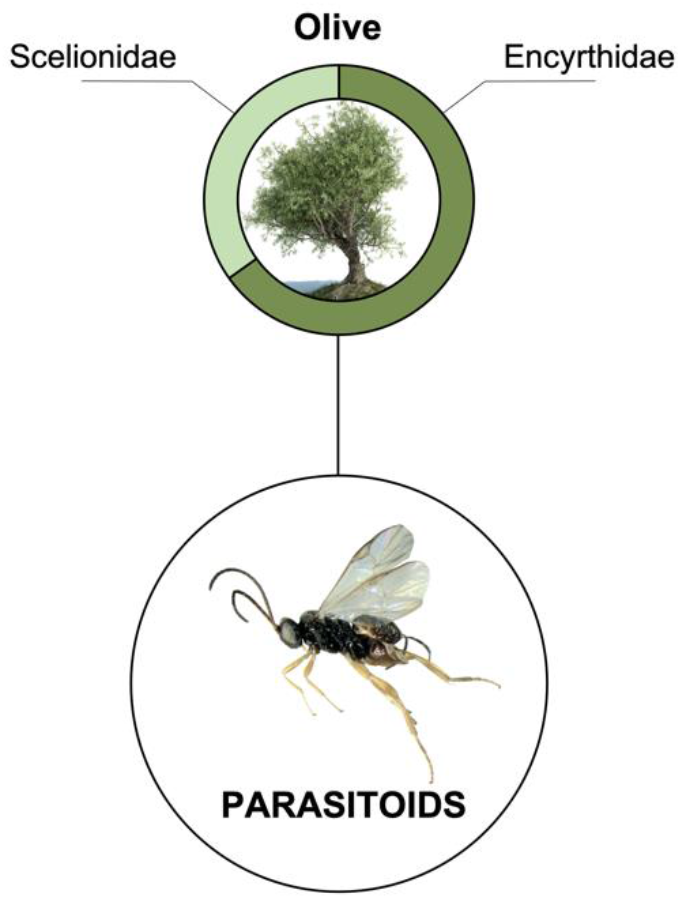
Families of parasitoids that had the highest effects in olive trees within the organic olive orchards. The circles show the proportion of the abundance amongst such families.

On the other hand, only two families were identified as omnivores: Formicidae (Hymenoptera) and Forficulidae (Dermaptera), the former had very high abundance and the latter almost none. Formicidae was present in all plants, tending to have more abundance in *R. sphaerocarpa*, *R. officinalis*, *T. zygis gracilis*, *L. longirrostris*, and the miscellaneous plants (Appendix).

Finally, only bees (Hymenoptera: Apidae) were identified as pollinators. Apidae was present in three plants within adjacent vegetation, four plants within ground cover, the miscellaneous plants and the olive trees, tending to have more abundance in *L. longirrostris* and olive trees (see Appendix).

### 3.2. Effects of habitat complexity

The abundance of the key families showed significant differences amongst study areas (*F*_4, 40_ = 2.708, *p* = 0.043), i.e., the areas with the highest complexity had the highest abundance (PI-2 and DEI: Tukey post hoc test, *p* = 0.037) (Fig. 3). Moreover, key family abundance had a positive relationship with the level of (plant) arrangement and scattering (*F*_1, 42_ = 10.661, *p* = 0.002) (Fig. 3) but not with the level of (plant) richness. On the other hand, the number of key families (number of taxa) did not follow any pattern, i.e., there is no difference amongst study areas (*F*_1, 43_ = 0.236, *p* = 0.915) and no relationship with the level of richness nor the level of arrangement and scattering.

**Fig. 3.**
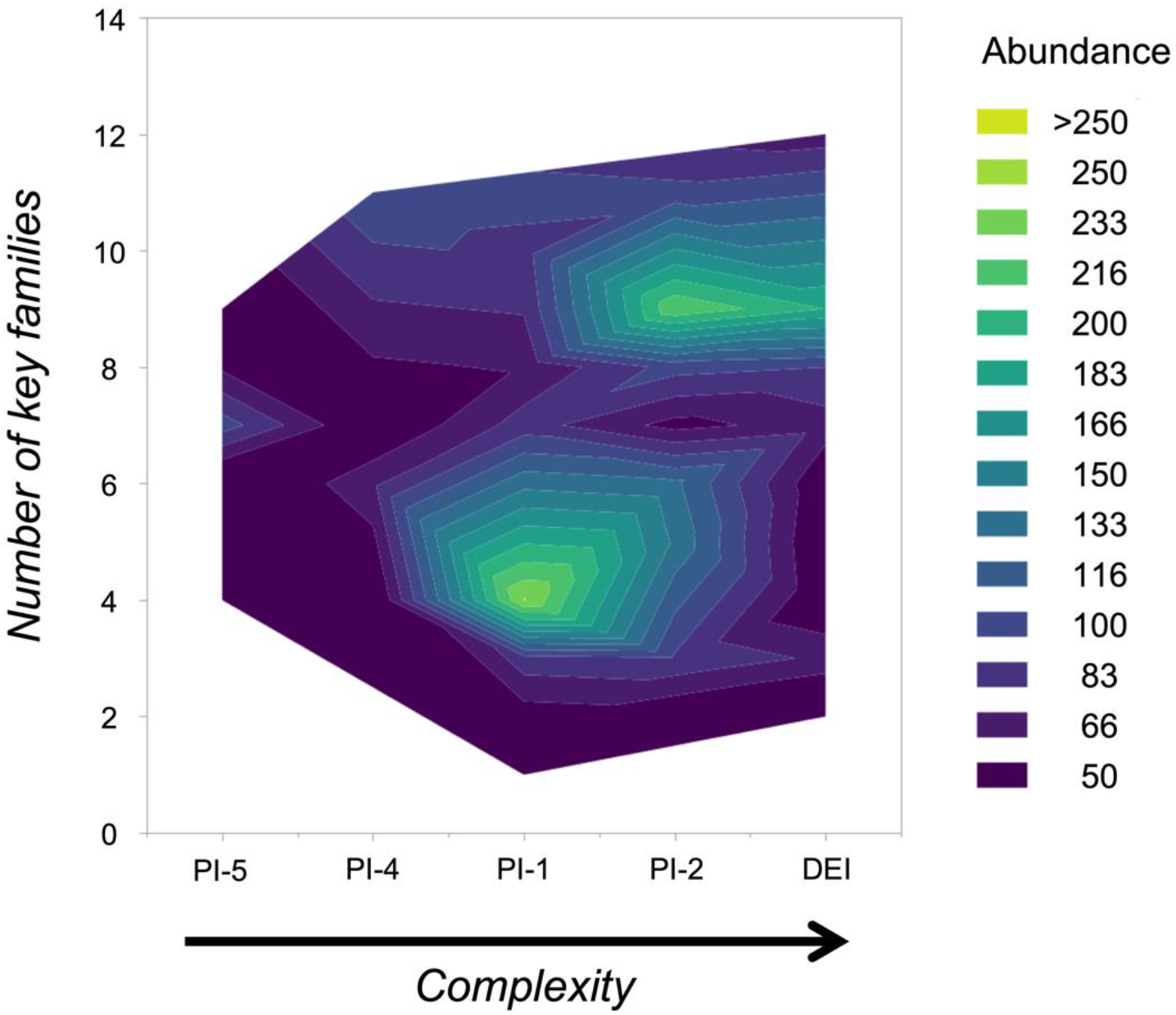
Contour plot showing the response values and desirable operating conditions. The contour plot contains the following elements: predictors on *X* (study areas arranged according to complexity) and *Y* (number of key families and abundance) axes. Contour lines connect points that have the same adjusted response value integrating data of total abundance of key families, i.e., the lines and its form is given by the proportion of the abundance amongst areas. The form of the aggregate is given by the number of families.

In addition, we found that the key families can be separated in four groups based on the form and topology of the smooth surfaces (Fig. 4). The NMDS showed that plant richness and plant arrangement and scattering affected differently the key families. For example, ants, ladybeetles, and uloborid spiders were mostly influenced by plant arrangement and scattering followed by the aeolothrips. Conversely, mirids and oxyopid and thomisid spiders were mostly influenced by plant richness followed by anthocorids, lacewings, parasitoids, and salticid spiders. Contrary to the anterior, the pollinator abundance was influenced by the less rich areas, where *L. longirrostris* had more presence. NMDS results suggest that the abundance and presence of each key family responds to the presence of specific plant species in each study area but key families form groups modulated by habitat complexity (primarily affected by the level of arrangement and scattering) (Fig. 4).

**Fig. 4.**
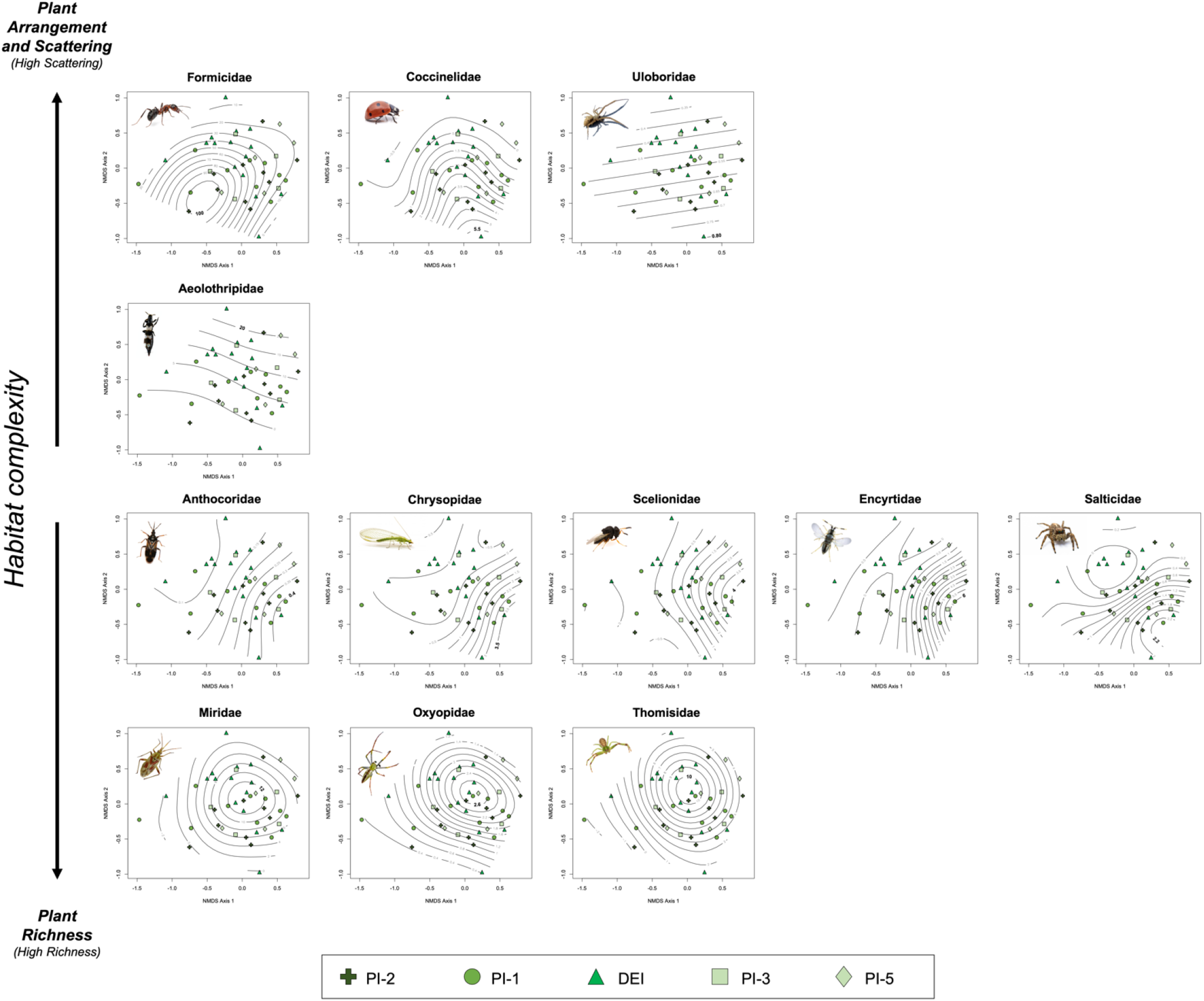
Non metric multi-dimensional scaling (NMDS) of the total abundance of natural enemies. Lines (smooth surfaces) represent different levels in the form of a gradient of each key family according to generalised additive models. Aggregates of families are given by the tendency in the smooth surface and arranged according to effects of plant richness or patch structure. Areas are arranged from most to least complex. To see details in each graph, proceed to the digital version of the figure in high definition.

## 4. Discussion

In this study we have assessed the effects that several plants within ground cover and adjacent vegetation may have to attract natural enemies and pollinators to olive orchards. We have also, assessed the effects of the components of habitat complexity on key families of natural enemies and pollinators.

Overall, from the 15 most representative plants in our study areas, only *A. radiatus*, *D. catholica*, and *L. longirrostris* showed effects to attract natural enemies and pollinators within ground cover and *G. cinerea speciosa*, *Q. rotundifolia*, *R. officinalis*, *T. zygis gracilis*, and *U. parviflorus* showed such an effect within adjacent vegetation.

Moreover, the miscellaneous plants also showed effects. This suggests that there are more plants in the adjacent vegetation that can attract beneficial arthropods compared with the ground cover. Perhaps, this is why in some modelling adjacent vegetation produced greater (positive) effects on predators than the ground cover, especially in shrubby habitats (Paredes et al., 2013; 2015). However, it has been shown that the ground cover maintains the highest abundance of natural enemies, i.e., parasitoids, predators, and omnivores together rather than the adjacent vegetation (Álvarez et al., 2019a), also it integrates more predators into the tropic network of the olive tree canopy (when ground cover is mature, Álvarez et al., 2019b), and promotes an effective predation of the olive moth *P. oleae* (Álvarez et al., 2021). Based on the type of beneficial arthropod, our results are in agreement with the findings of Álvarez et al., (2019a; 2021), Karamaouna et al., (2019), and Paredes et al., (2013). Accordingly, shrubby habitats had more attraction for spiders (predators) and ants (omnivores) but the ground cover had more attraction for bees (pollinators), heteropterans (predators), and members of aeolothripidae (predators). This supports the idea that adjacent vegetation and ground cover are different types of habitats, with different types of resources (and availability), which stay in synergy with the olive orchard. Nonetheless, habitat complexity plays an important role modulating the abundance of beneficial arthropods. Our results showed that abundance tend to increase as complexity increase when total abundance is measured (Fig. 3). Furthermore, it was interesting that only plant arrangement and scattering affected abundance rather than plant richness. This can be explained due to the fact that organic olive orchards are highly similar (e.g., management and structure). This is why there are no differences amongst our study areas when we use the level of richness, all areas are similar in their plant richness. Moreover, the richest areas turned to be the ones with higher abundance (Fig. 3) and according to NMDS analyses, richness do affect abundance specially the abundance of generalist and specialist predators, which is in agreement with theory (Haddad et al., 2001; Knops et al., 1999).

In regard to the anterior, Álvarez et al. (2019a) showed that the natural enemies of olive pests are more abundant in complex habitats and that they move across orchards and vegetations throughout the months of highest arthropod abundance. It seems that parasitoids, predators, and omnivores overwinter in the adjacent vegetation and when the temperature increases they move to the ground cover and, as a result of spill over, they can invade the olive trees predating new preys (such as olive pests). However, this movement is modulated only by the ground cover, i.e., predators and parasitoids invade the ground cover when it is growing, but when ground cover starts to wither the predators tend to return to the adjacent vegetation. Conversely, it is only at this time that parasitoids and omnivores move to the olive trees, which corresponds with the time that *P. oleae* lay their eggs on young olive fruit (Ramos et al., 1978). Accordingly, the synergy between adjacent vegetation and ground cover implies that complex habitats are paramount in order to increase natural enemy abundance as suggested by studies on natural ecosystems (Haddad et al., 2001; Knops et al., 1999), being the generalist arthropods the ones that will respond to such an effect. However, one has to take into account that the pattern showed here by abundance could be drove by few of the taxa present in one area and if it is wanted to increase the number of families or specific predators within olive orchards plant richness is of great importance as suggested by the NMDS (Fig. 4).

On the other hand, specialist predators of olive pests and parasitoids such as Anthocoridae, Chrysopidae (predators), Encyrtidae, and Scelionidae (parasitoids) showed effects for olive trees and tend to be (1) related with plant richness and (2) less affected by plant arrangement and scattering than other families (Fig. 4). For example, it is known that *Anthocoris nemoralis* (Anthocoridae) and *Chrysoperla carnea s.l.* (Chrysopidae) are important natural enemies that predate eggs and adults of *Prays oleae* (Morris et al., 1999; Villa et al., 2016b). In addition, the wasp *Ageniaspis fuscicollis* (Encyrtidae) is a polyembryonic parasitoid that lays its eggs on the eggs of *P. oleae* (Arambourg, 1984), and the wasp *Telenomus acrobater* (Scelionidae) is a hyperparasitoid that parasites the larvae of Chrysopidae (Campos, 1986). Hence, the specialist predators and parasitoids are linked to olive trees due to their need for specific prey, and thus, it is possible to enhance the abundance of such families by increasing habitat complexity but it would be paramount to increase the presence of the plant species in which these families have been recorded, aside from olive trees, within ground cover and adjacent vegetation (Alcalá-Herrera et al., 2019; Álvarez et al., 2021).

It is important to note that several of the natural enemies and the pollinators that have been recorded in this study had previously been known to inhabit the olive orchards and help to control olive pests (Torres, 2006). However, in our study Aeolothripidae showed a strong presence within the olive orchards. The role of the Aeolotrhipidae as natural enemies of olive pests has been poorly documented (reviewed by Torres, 2006). It is known that Aeolothripidae attack other Thysanoptera, but it has also been shown that some genera of Aeolothripidae can feed on mites, larvae, whiteflies, and aphids, as well as on the eggs of psyllids and lepidopterans (Lewis, 1973; Trdan et al., 2005). Their abundance suggests that they may be important assets amongst the natural enemies of olive pests as a parallel study suggests (Álvarez et al., 2021).

In regard to pollinators, we recorded the family Apidae as the only representative of this guild. Olive flowers are wind pollinated (Lavee, 1996) and insect pollination may supplement wind pollination. In the Mediterranean basin few insect pollinators of olive trees are known, the primary recorded groups are (1) bees of the families Apidae, Adrenidae, and Halictidae, and (2) hoverflies (Syrphidae), with the most representative being the honey bee *Apis mellifera* L. (Canale and Loni, 2010; Karamaouna et al., 2019). Apidae in this study were recorded in almost all the flowering plants, however, it seems that they are attracted to the ground cover because of the presence of yellowish flowers such as *L. longirrostris*, but they did visit the olive trees, which supports the results found by previous studies (Canale and Loni, 2010; Karamaouna et al., 2019).

## 5. Conclusions

Eight plants showed the best results regarding attracting beneficial arthropods within organic olive orchards. Key family abundance is affected by habitat complexity, i.e., the highest the complexity in a habitat the highest the abundances of natural enemies and pollinators, however, they are influenced differently by plant richness and plant arrangement and scattering. Our findings could be used by producers and technicians to increase the abundance of natural enemies and pollinators within olive orchards. We agree that these plant species have the potential to boost abundance in adjacent vegetation and ground cover, however high levels of complexity to conserve areas of natural habitats are paramount to produce such results, i.e., ground cover and adjacent vegetation must have several plant species within them, then they can be planned in order to integrate such plants according to the best features of arrangement and scattering.

## Acknowledgements

The authors want to thank Norberto Recio, Manuel Recio, and Rafael López Osorio who are the owners of the orchards; Marco Paganelli, Elena Torrente, Miguel Angel Sánchez and Rafael Cledera for their help identifying arthropods in the laboratory; and Raquel Jiménez and Carlos Martínez for their field assistance (the latter as part of an UGR collaboration grant). Special thanks to Antonio García and Jacinto Berzosa for their assistance identifying the plant species and the Thysanoptera, respectively and Mercedes Campos Arana for her support. H. A. Álvarez is grateful to Gemma Clemente Orta for her help conceptualizing graphic representations and CONACyT for providing him with an international PhD student grant (registry 332659). This study was financed by the Excellence Project of the Andalusian Regional Government (AGR 1419) obtained by Mercedes Campos Arana and F. Ruano.

## Appendix

Relative abundance (RA), trophic guild, months of record, and presence on plant species of all the families of arthropods (*n* = 106) identified in organic olive orchards. It is listed the plant species per type of vegetation: Adjacent vegetation, ground cover, and olive trees. The plants and months that had the highest records of abundance are showed in bold.

**Appendix.**

| Name |  | ID | RA (%) | Record on Type of vegetation: Plant | Month |
| --- | --- | --- | --- | --- | --- |
| Arachnida |  |  |  |  |  |
| Araneae | Amaurobiidae | Predator | 0,03 | ADJACENT: *U. parviflorus*. | June |
|  | Araneidae | Predator | 0,30 | ADJACENT: *Q. rotundifolia*, *R. officinalis*, *U. parviflorus*. G. COVER: *D. catholica*, *L. longirrostris*. OLIVE. | May, June, July |
|  | Dyctinidae | Predator | 0,02 | ADJACENT: *Q. rotundifolia*. | June, July |
|  | Linyphiidae | Predator | 0,14 | ADJACENT: *C. albidus*, *R. officinalis*, *T. zigys gracilis*. G. COVER: *A. radiatus*, *D. catholica*, *L. longirrostris*. | May, June |
|  | Mimetidae | Predator | 0,02 | G. COVER: *D. catholica*, miscellaneous. | May, June |
|  | Oxyopidae | Predator | 0,86 | ADJACENT: *C. albidus*, *R. officinalis*, *R. sphaerocarpa*, *T. mastichina*, *T. zigys gracilis*, *U. parviflorus*. G. COVER: *A. radiatus*, *L. longirrostris*, miscellaneous. OLIVE. | May, June, July |
|  | Philodromidae | Predator | 0,23 | ADJACENT: *C. albidus*, *G. cinerea speciosa*, *Q. rotundifolia*, *R. sphaerocarpa*, *U. parviflorus*. G. COVER: *D. catholica*, *R. officinalis*, *S. vulgaris*, miscellaneous. OLIVE. | May, June, July |
|  | Salticidae | Predator | 0,37 | ADJACENT: *P. dulcis*, *Q. rotundifolia*, *R. officinalis*, *R. sphaerocarpa*, *T. mastichina*, *T. zigys gracilis*, *U. parviflorus*. G. COVER: *L. longirrostris*, miscellaneous. OLIVE. | May, June, July |
|  | Sicariidae | Predator | 0,01 | ADJACENT: *U. parviflorus*. | June |
|  | Thomisidae | Predator | 2,88 | ADJACENT: *C. albidus*, *G. cinerea speciosa*, *P. dulcis*, *Q. rotundifolia*, *R. sphaerocarpa*, *R. officinalis*, *T. mastichina*, *T. zigys gracilis*, *U. parviflorus*. G. COVER: *A. radiatus*, *D. catholica*, *E. cicutarium*, *L. longirrostris*, *S. vulgaris*, miscellaneous. OLIVE. | May, June, July |
|  | Uoboridae | Predator | 0,27 | ADJACENT: *U. parviflorus*. | July |
|  | Zodariidae | Predator | 0,04 | ADJACENT: *U. parviflorus*. G. COVER: *L. longirrostris*. | June, July |
| Insecta |  |  |  |  |  |
| Blattodea | Blattellidae | Neutral arthropod | 0,09 | ADJACENT: *P. dulcis*, *Q. rotundifolia*, *R. officinalis*, *T. zigys gracilis*, *U. parviflorus*. | May, June, July |
| Coleoptera | Alleculidae | Neutral arthropod | 0,08 | ADJACENT: *C. albidus*, *R. sphaerocarpa*. G. COVER: *A. radiatus*. | May, June |
|  | Anthicidae | Neutral arthropod | 0,08 | ADJACENT: *T. zigys gracilis*. G. COVER: *L. longirrostris*, miscellaneous. | May, June, July |
|  | Apionidae | Neutral arthropod | 0,06 | ADJACENT: *C. albidus*. | June, July |
|  | Cantharidae | Predator | 0,03 | ADJACENT: *R. officinalis*, *T. zigys gracilis*. | May |
|  | Catopidae | Neutral arthropod | 0,09 | ADJACENT: *Q. rotundifolia*, *G. cinerea speciosa*, *R. sphaerocarpa*, *R. officinalis*. | May, June |
|  | Cerambycidae | Neutral arthropod | 0,91 | G. COVER: *L. longirrostris*. | May |
|  | Chrysomelidae | Neutral arthropod | 0,01 | ADJACENT: *C. albidus*, *P. dulcis*, *Q. rotundifolia*, *R. sphaerocarpa*, *R. officinalis*, *T. mastichina*, *T. zigys gracilis*, *U. parviflorus*. G. COVER: *D. catholica*, *L. longirrostris*, miscellaneous. OLIVE. | May, June, July |
|  | Cleridae | Predator | 0,04 | G. COVER: *L. longirrostris*. OLIVE. | May |
|  | Coccinellidae | Predator | 0,81 | ADJACENT: *C. albidus*, *P. dulcis*, *Q. rotundifolia*, *R. sphaerocarpa*, *R. officinalis*, *T. mastichina*, *T. zigys gracilis*, *U. parviflorus*. G. COVER: *A. radiatus*, *D. catholica*, *L. longirrostris*, miscellaneous. OLIVE. | May, June, July |
|  | Curculionidae | Neutral arthropod | 1,86 | ADJACENT: *C. albidus*, *G. cinerea speciosa*, *P. dulcis*, *Q. rotundifolia*, *R. sphaerocarpa*, *R. officinalis*, *T. mastichina*, *T. zigys gracilis*, *U. parviflorus*. G. COVER: *A. radiatus*, *D. catholica*, *E. cicutarium*, *L. longirrostris*, miscellaneous. OLIVE. | May, June, July |
|  | Dasytidae | Predator | 0,12 | ADJACENT: *R. sphaerocarpa*, *R. officinalis*, *T. zigys gracilis*, *U. parviflorus*. G. COVER: *D. catholica*, *L. longirrostris*. | May, June |
|  | Dermostidae | Neutral arthropod | 0,03 | ADJACENT: *Q. rotundifolia*. G. COVER: *D. catholica*, *L. longirrostris*. | May, June |
|  | Elateridae | Predator | 0,01 | G. COVER: *L. longirrostris*. | May |
|  | Malachiidae | Predator | 0,02 | ADJACENT: *U. parviflorus*. G. COVER: *L. longirrostris*. | May, June |
|  | Monotomidae | Neutral arthropod | 0,02 | G. COVER: miscellaneous. OLIVE. | June |
|  | Mycetophagidae | Neutral arthropod | 0,01 | ADJACENT: *Q. rotundifolia*. | May |
|  | Nitidulidae | Neutral arthropod | 0,03 | ADJACENT: *G. cinerea speciosa*. G. COVER: *A. radiatus*, *D. catholica*. | May |
|  | Phalacridae | Neutral arthropod | 0,61 | ADJACENT: *Q. rotundifolia*, *R. sphaerocarpa*, *R. officinalis*, *U. parviflorus*. G. COVER: *A. radiatus*, *L. longirrostris*. OLIVE. | May, June, July |
|  | Ptinidae | Neutral arthropod | 0,01 | ADJACENT: *P. dulcis*. | June |
|  | Scarabaeidae | Neutral arthropod | 0,01 | G. COVER: *D. catholica*. | May |
|  | Staphylinidae | Predator | 0,04 | ADJACENT: *R. officinalis*. G. COVER: *A. radiatus*, *L. longirrostris*. | May |
| Dermaptera | Forficulidae | Omnivore | 0,02 | ADJACENT: *P. dulcis*, *G. cinerea speciosa*. | May, June |
| Diptera | Agromyzidae | Neutral arthropod | 0,02 | OLIVE. | June, July |
|  | Asilidae | Predator | 0,01 | ADJACENT: *Q. rotundifolia*. | May |
|  | Bibionidae | Neutral arthropod | 0,03 | ADJACENT: *Q. rotundifolia*, *R. officinalis*. OLIVE. | May, July |
|  | Bombyliidae | Neutral arthropod | 0,09 | ADJACENT: *R. sphaerocarpa*, *R. officinalis*, *T. mastichina*, *U. parviflorus*. G. COVER: *L. longirrostris*. OLIVE. | May, June, July |
|  | Calliphoridae | Neutral arthropod | 0,13 | ADJACENT: *Q. rotundifolia*. | June |
|  | Camillidae | Neutral arthropod | 0,01 | G. COVER: *D. catholica*, *E. cicutarium*, *L. longirrostris*, *miscellaneous*. | **May**, June |
|  | Cecidomyiidae | Neutral arthropod | 0,36 | ADJACENT: *G. cinerea speciosa*, *R. officinalis*, *T. mastichina*, *T. zigys gracilis*, *U. parviflorus*. G. COVER: *D. catholica*, *L. longirrostris*, **miscellaneous**. | May, **June**, July |
|  | Ceratopogonidae | Neutral arthropod | 0,01 | ADJACENT: *T. zigys gracilis*. | June |
|  | Chamaemyiidae | Predator | 0,01 | ADJACENT: *T. zigys gracilis*. | June |
|  | Chironomidae | Neutral arthropod | 0,06 | ADJACENT: *Q. rotundifolia*. G. COVER: *miscellaneous*. OLIVE. | May, **June** |
|  | Chloropidae | Neutral arthropod | 0,22 | ADJACENT: *P. dulcis*, *R. sphaerocarpa*, *R. officinalis*. G. COVER: *C. melitenses*, *L. longirrostris*, *miscellaneous*. OLIVE. | May, June, **July** |
|  | Dolichopodidae | Predator | 0,16 | ADJACENT: *P. dulcis*, *Q. rotundifolia*, *T. zigys gracilis*. G. COVER: *C. melitenses*, *L. longirrostris*, *miscellaneous*. | **May**, June, July |
|  | Empididae | Neutral arthropod | 0,08 | ADJACENT: *P. dulcis*, *Q. rotundifolia*, *T. zigys gracilis*. OLIVE. | **May**, June |
|  | Lauxaniidae | Neutral arthropod | 0,01 | G. COVER: *miscellaneous*. | June |
|  | Limonidae | Neutral arthropod | 0,02 | ADJACENT: *P. dulcis*. | May, June |
|  | Muscidae | Neutral arthropod | 0,05 | ADJACENT: *Q. rotundifolia*. G. COVER: *D. catholica*. OLIVE. | May, June, July |
|  | Mythicomyiidae | Neutral arthropod | 0,05 | G. COVER: *miscellaneous*. | July |
|  | Opomyzidae | Neutral arthropod | 0,01 | G. COVER: *D. catholica*. | May |
|  | Phoridae | Neutral arthropod | 0,01 | ADJACENT: *Q. rotundifolia*. | May |
|  | Sciaridae | Neutral arthropod | 0,12 | ADJACENT: *R. officinalis*, *U. parviflorus*. G. COVER: *D. catholica*, *E. cicutarium*, **miscellaneous**. | **May**, June, July |
|  | Tephritidae | Neutral arthropod | 0,22 | ADJACENT: *P. dulcis*, *Q. rotundifolia*, *R. officinalis*, *U. parviflorus*. G. COVER: *D. catholica*, *L. longirrostris*, *S. vulgaris*. OLIVE. | May, **June** |
|  | Heleomyzidae | Neutral arthropod | 0,04 | G. COVER: *D. catholica*, *miscellaneous*. | May, June |
| Heteroptera | Anthocoridae | Predator | 0,11 | ADJACENT: *C. albidus*, *T. zigys gracilis*. G. COVER: *miscellaneous*. **OLIVE**. | May, **June** |
|  | Aphididae | Neutral arthropod | 21,11 | ADJACENT: *P. dulcis*, *Q. rotundifolia*, *R. sphaerocarpa*, *R. officinalis*, *T. matchina*, *T. zigys gracilis*, *U. parviflorus*. G. COVER: *A. radiatus*, *D. catholica*, *E. cicutarium*, *L. longirrostris*, *S. vulgaris*, **miscellaneous**. OLIVE. | May, **June**, July |
|  | Berytidae | Neutral arthropod | 0,20 | ADJACENT: *T. zigys gracilis*, *U. parviflorus*. G. COVER: *A. radiatus*, *L. longirrostris*, *S. vulgaris*, **miscellaneous**. | May, **June**, July |
|  | Coccidae | Neutral arthropod | 0,28 | ADJACENT: *C. albidus*, *T. zigys gracilis*, *U. parviflorus*. G. COVER: *D. catholica*, *miscellaneous*. **OLIVE**. | May, **June**, July |
|  | Coreidae | Neutral arthropod | 0,01 | G. COVER: *miscellaneous*. | June |
|  | Cydnidae | Neutral arthropod | 0,01 | ADJACENT: *T. zigys gracilis*. | May |
|  | Fulgoromorpha | Neutral arthropod | 13,56 | ADJACENT: *C. albidus*, *G. cinerea speciosa*, *P. dulcis*, *Q. rotundifolia*, *R. sphaerocarpa*, *R. officinalis*, *T. matchina*, *T. zigys gracilis*, *U. parviflorus*. G. COVER: *A. radiatus*, *C. melitenses*, *D. catholica*, *E. cicutarium*, *L. longirrostris*, *S. vulgaris*, *miscellaneous*. **OLIVE**. | **May**, June, July |
|  | Geocoridae | Predator | 0,02 | ADJACENT: *U. parviflorus*. | July |
|  | Lygaeidae | Neutral and predator (facultative) | 0,56 | ADJACENT: *C. albidus*, *R. officinalis*, *T. mastichina*, *T. zigys gracilis*, *U. parviflorus*. G. COVER: *C. melitenses*, *A. radiatus*, *L. longirrostris*, *S. vulgaris*, **miscellaneous**. OLIVE. | May, **June**, July |
|  | Miridae | Predator | 3,55 | ADJACENT: *C. albidus*, *G. cinerea speciosa*, *Q. rotundifolia*, *R. sphaerocarpa*, *R. officinalis*, *T. mastichina*, *T. zigys gracilis*, *U. parviflorus*. G. COVER: *A. radiatus*, *C. melitenses*, *D. catholica*, *E. cicutarium*, *L. longirrostris*, *S. vulgaris*, *miscellaneous*. **OLIVE**. | May, **June**, July |
|  | Nabidae | Predator | 0,03 | ADJACENT: *P. dulcis*, *U. parviflorus*. G. COVER: *miscellaneous*. | May, June |
|  | Pentatomidae | Neutral arthropod | 0,53 | ADJACENT: *G. cinerea speciosa*, *R. sphaerocarpa*, *R. officinalis*, *T. mastichina*, *T. zigys gracilis*, *U. parviflorus*. G. COVER: *A. radiatus*, *D. catholica*, *L. longirrostris*, *miscellaneous*. | **May**, June, July |
|  | Plataspidae | Neutral arthropod | 0,01 | G. COVER: *miscellaneous*. | July |
|  | Psyllidae | Pest | 12,52 | ADJACENT: *C. albidus*, *G. cinerea speciosa*, *Q. rotundifolia*, *R. sphaerocarpa*, *R. officinalis*, *T. zigys gracilis*, *U. parviflorus*. G. COVER: *C. melitenses*, *L. longirrostris*, *miscellaneous*. **OLIVE**. | May, June, **July** |
|  | Rhopalidae | Neutral arthropod | 0,32 | ADJACENT: *Q. rotundifolia*, *R. officinalis*, *T. zigys gracilis*, *U. parviflorus*. G. COVER: *A. radiatus*, *E. cicutarium*, *S. vulgaris*, *miscellaneous*. OLIVE. | May, **June**, July |
|  | Tingidae | Neutral arthropod | 0,20 | ADJACENT: *P. dulcis*, *Q. rotundifolia*, *R. officinalis*, *T. zigys gracilis*, *U. parviflorus*. G. COVER: *L. longirrostris*. | **May**, June, July |
| Hymenoptera | Aphelinidae | Neutral arthropod | 0,05 | OLIVE. | June |
|  | Apidae | Pollinator | 0,51 | ADJACENT: *C. albidus*, *Q. rotundifolia*, *R. officinalis*. G. COVER: *A. radiatus*, *D. catholica*, **L. longirrostris**, *S. vulgaris*, miscellaneous. **OLIVE**. | **May**, June, July |
|  | Bethylidae | Parasitoid | 0,14 | ADJACENT: *R. officinalis*. G. COVER: **L. longirrostris**. OLIVE. | May |
|  | Braconidae | Parasitoid | 0,45 | ADJACENT: *P. dulcis*, *Q. rotundifolia*, *U. parviflorus*. G. COVER: *A. radiatus*, **D. catholica**, **L. longirrostris**, *S. vulgaris*, miscellaneous. OLIVE. | **May**, June |
|  | Ceraphronidae | Neutral arthropod | 0,01 | ADJACENT: *R. officinalis*. | May |
|  | Chrysididae | Parasitoid | 0,02 | ADJACENT: *Q. rotundifolia*. OLIVE. | May |
|  | Cynipidae | Neutral arthropod | 0,02 | ADJACENT: *Q. rotundifolia*, *U. parviflorus*. | June |
|  | Diapriidae | Parasitoid | 0,02 | G. COVER: *D. catholica*, *S. vulgaris*. | May, June |
|  | Elasmidae | Parasitoid | 0,10 | ADJACENT: *Q. rotundifolia*, *T. mastichina*. G. COVER: miscellaneous. OLIVE. | May, **June** |
|  | Encyrtidae | Parasitoid | 0,83 | ADJACENT: **Q. rotundifolia**, *R. officinalis*, *T. zigys gracilis*, *U. parviflorus*. G. COVER: *D. catholica*, *L. longirrostris*, miscellaneous. **OLIVE**. | May, **June**, July |
|  | Eulophidae | Parasitoid | 0,13 | ADJACENT: *P. dulcis*, *R. officinalis*, *T. mastichina*, *T. zigys gracilis*. G. COVER: **D. catholica**, miscellaneous. | **May**, June |
|  | Eupelmidae | Parasitoid | 0,02 | ADJACENT: *T. zigys gracilis*. | June |
|  | Eurytomidae | Parasitoid | 0,03 | ADJACENT: *P. dulcis*, *Q. rotundifolia*. | May, June |
|  | Figitidae | Parasitoid | 0,01 | G. COVER: miscellaneous. | June |
|  | Formicidae | Omnivore | 21,55 | ADJACENT: *C. albidus*, *G. cinerea speciosa*, *Q. rotundifolia*, *P. dulcis*, **R. sphaerocarpa**, **R. officinalis**, *T. mastichina*, **T. zigys gracilis**, *U. parviflorus*. G. COVER: *A. radiatus*, *C. melitenses*, *D. catholica*, *E. cicutarium*, **L. longirrostris**, *S. vulgaris*, **miscellaneous**. OLIVE. | **May**, **June**, July |
|  | Ichneumonidae | Parasitoid | 0,04 | ADJACENT: *Q. rotundifolia*, *R. sphaerocarpa*. G. COVER: *L. longirrostris*. | **May**, June |
|  | Leucospidae | Neutral arthropod | 0,01 | ADJACENT: *Q. rotundifolia*. | June |
|  | Megaspilidae | Parasitoid | 0,01 | ADJACENT: *Q. rotundifolia*. | May |
|  | Mymaridae | Parasitoid | 0,09 | ADJACENT: *R. officinalis*. G. COVER: *S. vulgaris*, miscellaneous. OLIVE. | May, **June** |
|  | Platygastridae | Parasitoid | 0,04 | ADJACENT: *G. cinerea speciosa*, *Q. rotundifolia*. G. COVER: *L. longirrostris*. | May, June |
|  | Pompilidae | Neutral arthropod | 0,02 | ADJACENT: *P. dulcis*, *Q. rotundifolia*. | June |
|  | Pteromalidae | Parasitoid | 0,45 | ADJACENT: *C. albidus*, *G. cinerea speciosa*, *P. dulcis*, *Q. rotundifolia*, *R. officinalis*, *T. mastichina*, *T. zigys gracilis*, *U. parviflorus*. G. COVER: **D. catholica**, **L. longirrostris**, *S. vulgaris*, **miscellaneous**. OLIVE. | May, **June**, July |
|  | Scelionidae | Parasitoid | 0,63 | ADJACENT: *G. cinerea speciosa*, *Q. rotundifolia*, *T. mastichina*, *T. zigys gracilis*, *U. parviflorus*. G. COVER: **D. catholica**, **L. longirrostris**, *S. vulgaris*, **miscellaneous**. **OLIVE**. | May, **June**, July |
|  | Sphecidae | Parasitoid and predator | 0,01 | ADJACENT: *R. officinalis*. | May |
| Lepidoptera | Praydidae | Pest | 0,33 | ADJACENT: *R. officinalis*. G. COVER: miscellaneous. **OLIVE**. | May, **June** |
| Mantodea | Mantidae | Predator | 0,08 | ADJACENT: *P. dulcis*, *Q. rotundifolia*, *R. officinalis*, *T. zigys gracilis*, *U. parviflorus*. G. COVER: miscellaneous. | **June**, July |
| Neuroptera | Chrysopidae | Predator | 0,37 | ADJACENT: *C. albidus*, **Q. rotundifolia**, *P. dulcis*, *T. zigys gracilis*. G. COVER: *D. catholica*, miscellaneous. **OLIVE**. | May, **June**, July |
|  | Coniopterygidae | Predator | 0,03 | ADJACENT: *Q. rotundifolia*. | May, June |
| Phasmida | Phasmatidae | Neutral arthropod | 0,01 | ADJACENT: *T. zigys gracilis*. | June |
| Raphidioptera | Raphidiidae | Predator | 0,02 | ADJACENT: *Q. rotundifolia*, *R. sphaerocarpa*. | May, June |
| Thysanoptera | Aeolothripidae | Predator | 3,47 | ADJACENT: *G. cinerea speciosa*, *Q. rotundifolia*, *R. officinalis*, *T. zigys gracilis*, *U. parviflorus*. G. COVER: *A. radiatus*, **D. catholica**, *E. cicutarium*, **L. longirrostris**, miscellaneous. OLIVE. | **May**, June, July |
|  | Phlaeothripidae | Neutral arthropod | 1,39 | ADJACENT: *G. cinerea speciosa*, *Q. rotundifolia*, *R. sphaerocarpa*, *T. mastichina*, **T. zigys gracilis**, *U. parviflorus*. G. COVER: *A. radiatus*, *C. melitenses*, *D. catholica*, *L. longirrostris*, miscellaneous. OLIVE. | **May**, June, July |
|  | Thripidae | Neutral arthropod | 4,16 | ADJACENT: *Q. rotundifolia*, **R. sphaerocarpa**, *R. officinalis*, *T. zigys gracilis*, *U. parviflorus*. G. COVER: *C. melitenses*, *D. catholica*, *E. cicutarium*, **L. longirrostris**, *S. vulgaris*, **miscellaneous**. OLIVE. | May, **June**, July |

## References

Alcalá-Herrera, R., Ruano, F., Ramírez, C.G., Frischie, S., Campos, M., 2019. Attraction of green lacewings (Neuroptera: Chrysopidae) to native plants used as ground cover in woody Mediterranean agroecosystems. Biol. Control 139, 104066.

Álvarez, H.A., Carrillo-Ruiz, H., Morón, M.A., 2016. Record of Scarabaeoidea larvae and adults associated with *Amaranthus hypochondriacus* L. and living fences. Southwest. Entomol. 41, 675–679.

Álvarez, H.A., Carrillo-Ruiz, H., Jiménez-García, D., Morón, M.A., 2017. Abundance of insect fauna associated with *Amaranthus hypochondriacus* L. Crop, in relation to natural living fences. Southwest. Entomol. 42, 131–135.

Álvarez, H.A., Morente, M., Oi, F.S., Rodríguez, E., Campos, M., Ruano, F., 2019a. Semi-natural habitat complexity affects abundance and movement of natural enemies in organic olive orchards. Agric. Ecosyst. Environ. 285, 106618.

Álvarez, H.A., Morente, M., Campos, M., Ruano, F., 2019b. La madurez de las cubiertas vegetales aumenta la presencia de enemigos naturales y la resiliencia de la red trófica de la copa del olivo. Ecosistemas 28, 92–106.

Álvarez, H. A., Jiménez-Muñoz, R., Morente, M., Campos, M., Ruano, F., 2021. Ground cover presence in organic olive orchards affects the interaction of natural enemies against *Prays oleae*, promoting an effective egg predation. Biorxiv. doi: https://doi.org/10.1101/2021.02.03.429537

Arambourg, Y., 1986. Traité d’Entomologie Oleicole. Consejo Oleícola Internacional, Madrid, Spain, 360 p.

Balmford, A., Green, R., Phalan, B., 2012. What conservationists need to know about farming. Proc. R. Soc. B rspb20120515.

Campos, M., 1986. Influencia del complejo parasitario sobre las poblaciones de *Chrysoperla carnea* (Neuroptera, Chrysopidae) en olivares del sur de España. Neuroptera Int. 4, 97–105.

Canale, A., Loni, A., 2010. Insects visiting olive flowers (*Olea europaea* L.) in a Tuscan olive grove. Redia 92, 95–98.

Clemente-Orta, G., Álvarez, H.A., 2019. La influencia del paisaje agrícola en el control biológico desde una perspectiva espacial. Ecosistemas 28, 13–25.

Clemente-Orta, G., Madeira, F., Batuecas, I., Sossai, S., Juárez-Escario, A., Albajes, R., 2020. Changes in landscape composition influence the abundance of insects on maize: The role of fruit orchards and alfalfa crops. Agric. Ecosyst. Environ. 291, 106805.

Junta de Andalucía, 2007. 5.4. Áreas Forestales y Naturales. In: Guía técnica del Mapa de Usos y Coberturas Vegetales del Suelo de Andalucía 1:25.000. Junta de Andalucía. Consejería de Agricultura, Ganadería, Pesca y Desarrollo Sostenible (ámbito de Desarrollo Sostenible). p 148–202. http://www.juntadeandalucia.es/medioambiente/site/rediam/menuitem.04dc44281e5d53cf8ca78ca731525ea0/?vgnextoid=de07cb4af9245110VgnVCM1000000624e50aRCRD

Gkisakis, V., Volakakis, N., ollaros, D., Brberi, P., Kabourakis, E.M., 2016. Soil arthropod community in the olive agroecosystem: determined by environment and farming practices in different management systems and agroecological zones. Agric. Ecosyst. Environ. 218, 178–189.

Haddad, N.M., Tilman, D., Haarstad, J., Ritchie, M., Knops, J.M., 2001. Contrasting effects of plant richness and composition on insect communities: a field experiment. Am. Nat. 158, 17–35.

Hatt, S., Uyttenbroeck, R., Lopes, T., Mouchon, P., Chen, J.L., Piqueray, J., Monty, A., Francis, F., 2017. Do flower mixtures with high functional diversity enhance aphid predators in wildflower strips? Eur. J. Entomol. 114, 66–76.

Karamaouna, F., et al., 2019. Ground cover management with mixtures of flowering plants to enhance insect pollinators and natural enemies of pests in olive groves. Agric. Ecosyst. Environ. 274, 76–89.

Karp, D.S., et al., 2018. Crop pests and predators exhibit inconsistent responses to surrounding landscape composition. Proc. Nat. Acad. Sci. 115, E7863–E7870.

Knops, J.M., et al., 1999. Effects of plant species richness on invasion dynamics, disease outbreaks, insect abundances and diversity. Ecol. Lett. 2, 286–293.

Laurance, W.F., 2007. Ecosystem decay of Amazonian forest fragments: implications for conservation. In: Tscharntke, T., Leuschner, C., Zeller, M., Guhardja, E., Bidin, A. (Eds.), Stability of Tropical Rainforest Margins. Springer, Berlin, pp. 9–35.

Lavee, S., 1996. Biologia e fisiologia dell’olivo. In: Enciclopedia Mondiale dell’Olivo, Consejo Oleícola Internacional, Madrid, pp.61–110.

Lewis, T. 1973. Thrips – their biology, ecology and economic importance. Academic Press, London & New York. 349 p.

López-Barrera, F., Armesto, J.J., Williams-Linera, G., Smith-Ramírez, C., Manson, R.H., 2007. Fragmentation and edge effects on plant–animal interactions, ecological processes and biodiversity. In: Newton, A.C. (Ed.), Biodiversity Loss and Conservation in Fragmented Forest Landscapes. The Forests of Montane Mexico and Temperate South America. CABI, Wallingford, Oxfordshire, UK, pp. 69–101.

Malavolta, C., Perdikis, D. 2018. Crop Specific Technical Guidelines for Integrated Production of Olives. IOBC-WPRS Commission IP Guidelines. https://www.iobc-wprs.org/members/shop_en.cfm?mod_Shop_detail_produkte=193

Morris, T.I., Campos, M., Kidd, N.A.C., Symondson, W.O.C., 1999. What is consuming *Prays oleae* (Bernard) (Lep.: Yponomeutidae) and when: a serological solution?. Crop Prot. 18, 17–22.

Nave, A., Gonçalves, F., Crespí, A.L., Campos, M., Torres, L., 2016. Evaluation of native plant flower characteristics for conservation biological control of *Prays oleae*. Bull. Entomol. Res. 106, 249–257.

Oldfield, S., 2019. The US National Seed Strategy for Rehabilitation and Restoration: progress and prospects. Plant Biol. 21, 380–382.

Paredes, D., Cayuela, L., Campos, M., 2013. Synergistic effects of ground cover and adjacent vegetation on natural enemies of olive insect pests. Agric. Ecosyst. Environ. 173, 72–80.

Paredes, D., Cayuela, L., Gurr, G.M., Campos, M., 2015. Is ground cover vegetation an effective biological control enhancement strategy against olive pests?. PLoS One 10, e0117265.

Pedrini, S., Lewandrowski, W., Stevens, J.C., Dixon, K.W., 2019. Optimising seed processing techniques to improve germination and sowability of native grasses for ecological restoration. Plant Biol. 21, 415–424.

Porcel, M., Cotes, B., Castro, J., Campos, M., 2017. The effect of resident vegetation cover on abundance and diversity of green lacewings (Neuroptera: Chrysopidae) on olive trees. J. Pest Sci. 90, 195–206.

R Development Core Team, 2019. R: a Language and Environment for Statistical Computing. Available at Vienna: R Foundation for Statistical Computing. http://www.rproject.org/.

Ramos, P., Campos, M., Ramos, J.M., 1978. Factores limitantes en la fluctuación de poblaciones de *Prays oleae* Bern. Bol. San. Veg. Plagas 4, 1–6.

Ruano, F., Lozano, C., García, P., Pena, A., Tinaut, A., Pascual, F., Campos, M., 2004. Use of arthropods for the evaluation of the olive-orchard management regimes. Agr. Forest Entomol. 6, 111–120.

Rusch, A., Valantin-Morison, M., Sarthou, J.P., Roger-Estrade, J., 2010. Biological control of insect pests in agroecosystems: effects of crop management, farming systems, and seminatural habitats at the landscape scale: a review. Adv. Agron. 109, 219–259.

Santos, S.A.P., Pereira, J.A., Torres, L.M., Nogueira, A.J.A., 2007. Evaluation of the effects, on canopy arthropods, of two agricultural management systems to control pests in olive groves from north-east of Portugal. Chemosphere 67, 131–139. https://doi.org/10.1016/j.chemosphere.2006.09.014.

Trdan, S., Andjus, L., Raspudić, E., Kač, M., 2005. Distribution of *Aeolothrips intermedius* Bagnall (Thysanoptera: Aeolothripidae) and its potential prey Thysanoptera species on different cultivated host plants. J. Pest Sci. 78, 217–226.

Torres, L., 2006. A fauna auxiliar do olival ea sua conservação. Joao Azevedo, 92 p.

Tscharntke, T., et al., 2016. When natural habitat fails to enhance biological pest control Five hypotheses. Biol. Control 204, 449–458.

Ver Hoef, J.M., Boveng, P.L., 2007. Quasi-poisson vs. Negative binomial regression: how should We model overdispersed count data? Ecology 88, 2766–2772.

Van Rijn, P.C.J., Wackers, F.L., 2016. Nectar accessibility determines fitness, flower choice and abundance of hoverflies that provide natural pest control. J. Appl. Ecol. 53, 925–933.

Villa, M., Santos, S.A., Mexia, A., Bento, A., Pereira, J.A., 2016a. Ground cover management affects parasitism of *Prays oleae* (Bernard). Biol. Control 96, 72–77.

Villa, M., Santos, S.A., Benhadi-Marín, J., Mexia, A., Bento, A., Pereira, J.A., 2016b. Life-history parameters of *Chrysoperla carnea sl* fed on spontaneous plant species and insect honeydews: importance for conservation biological control. BioControl 61, 533–543.

Villa, M., Marrão, R., Mexia, A., Bento, A., Pereira, J.A., 2016c. Are wild flowers and insect honeydews potential food resources for adults of the olive moth, *Prays oleae*?. J. Pest Sci. 90, 185–194.

Wan, N.F., Gu, X.J., Ji, X.Y., Jiang, J.X., Wu, J.H., Li, B., 2014. Ecological engineering of ground cover vegetation enhances the diversity and stability of peach orchard canopy arthropod communities. Ecol. Eng. 70, 175–182.

Wan, N.F., et al., 2018. Increasing plant diversity with border crops reduces insecticide use and increases crop yield in urban agriculture. eLife 7, e35103.

